# Limited transgenerational effects of environmental temperatures on thermal performance of a cold-adapted salmonid

**DOI:** 10.1101/2020.05.13.094318

**Authors:** Chantelle M. Penney, Gary Burness, Joshua Robertson, Chris C. Wilson

## Abstract

The capacity of ectotherms to cope with rising temperatures associated with climate change is a significant conservation concern as the rate of warming is likely too fast for adaptation to occur in some populations. Transgenerational plasticity, if present, could potentially buffer some of the negative impacts of warming on future generations. We examined transgenerational plasticity in lake trout to assess their inter-generational potential to cope with anticipated warming. We acclimated adult lake trout to cold or warm temperatures for several months, then bred them to produce offspring from parents of matched and mismatched temperatures. At the fry stage, offspring were also acclimated to cold or warm temperatures and their thermal performance was assessed by measuring their critical thermal maximum and metabolic rate during an acute temperature challenge. Overall, transgenerational plasticity was evident: thermal performance of offspring reflected both maternal and paternal environmental conditions, and offspring performed better when their environment matched that of their parents. There was little variation in offspring critical thermal maximum or peak metabolic rate, although cold-acclimated offspring from warm-acclimated parents exhibited elevated standard metabolic rates, suggesting that transgenerational effects can be detrimental when parent and offspring environments mismatch. These results demonstrate both the occurrence and limitations of transgenerational plasticity in a coldwater salmonid in response to elevated temperature, as well as potential ecological risks associated with transgenerational plasticity when an environmental change experienced by the parents does not persist with the next generation.

## Introduction

Populations are being forced to respond to climate change as environmental temperatures continue to increase towards viable limits (Hazen *et al*., 2013; Galbraith *et al*., 2014; Luo *et al*., 2015). Many species are resorting to migration and range shifts where movement to more suitable habitats is possible (e.g. freshwater fish: Chu *et al*., 2005; birds: VanDerWal *et al*., 2013; mussels: Inoue *et al*., 2017), but those that are unable to relocate will need to acclimatize or adapt to warmer conditions if they are to persist. Phenotypic, developmental, and transgenerational plasticity along with genetic changes can influence adaptation of populations to climate change (Bernatchez, 2016; Smith *et al*., 2016). Organisms with little phenotypic plasticity may not be able to acclimatize to projected climatic conditions (Somero, 2010; Kelly *et al*., 2014) and adaptation may also not be feasible for populations that are small, isolated, or have adapted to thermally stable habitats because they are expected to have reduced standing genetic variation and less evolutionary potential (Willi *et al*., 2006; Meier *et al*., 2014). For species with long generation times, the rate of environmental change may also outpace the spread of beneficial genes (O’Grady *et al*., 2008; Visser, 2008; Munday *et al*., 2013).

Populations may be able to compensate for long-term changes in temperature by preconditioning their offspring for harsher environments (Yin *et al*., 2019) which may, over time, influence adaptation (Bonduriansky *et al*., 2012). This preconditioning can involve maternal and paternal (non-genetic) effects including nutrient provisioning of the eggs, transfer of hormones and other cytoplasmic components, and inheritance of epigenetic factors which can change the way genes are expressed (Deans & Maggert, 2015; Charlesworth *et al*., 2017). Epigenetic factors include modifications in histone configuration, DNA methylation, expression of microRNA or changes in chromatin structure (Hanson & Skinner, 2016), each of which can be induced by changes in the environment. This non-genetic inheritance can be observed through studies of transgenerational plasticity (TGP), which is a plastic response that occurs when the effects of the parent’s environment appear in the offspring’s phenotype (Bell & Hellman, 2019). TGP has been documented in several fish species including three-spined stickleback (Shama et al. 2014), sheepshead minnow (Salinas & Munch 2012), and tropical damselfish (Donelson *et al*., 2012; Munday *et al*., 2017). These fish are warm-adapted or eurythermal species, and it has not yet been confirmed whether this response also occurs in cold-adapted, stenothermal ectotherms. It is also unclear whether TGP is contingent on existing genetic variation which is relevant to populations that have adapted to cold, stable environments since they are likely to have experienced genomic loss over time (Willi et al,. 2006; Wilson, 2017).

The lake trout (*Salvelinus namaycush*) is a cold-adapted, stenothermal salmonid (Martin & Olver, 1980; Casselman, 2008) under significant threat from climate change (Evans, 2007; Guzzo & Blanchfield, 2017). These fish are restricted to oligotrophic lakes, preferring temperatures between 10-12 °C (Edsall, 2000; Martinez *et al*., 2009), and their habitat is transforming due to climate change: lake surface temperatures are increasing, the length of time that lakes are covered by ice is shortening, and the extent of cool, highly-oxygenated refuges are becoming limited during the summer (Reist *et al*., 2016; Guzzo & Blanchfield, 2017). These environmental changes already have an observable negative impact on lake trout as warmer temperatures at spawning reduces the survival of the fry at hatch (Casselman *et al.*, 2002). Furthermore, evidence suggests that genetic variation is low for some populations of lake trout (Perrier *et al*., 2017), and there is little variation in the capacity for within-generation temperature acclimation among allopatric populations (Kelly *et al*., 2014). These attributes make the lake trout a fitting species to study TGP in cold-adapted, stenothermal organisms. Their limited within-generation plasticity provides an opportunity to understand how TGP fits within the scope of possible thermal responses of organisms that are forced to cope with climate change.

We hypothesized that transgenerational plasticity occurs in cold-adapted, stenothermal ectotherms, potentially enabling them to cope with warmer environments. To test this hypothesis, we acclimated hatchery-raised, adult lake trout to cool (optimal) and warm temperatures, then used a full factorial mating design to cross fish within and between temperatures. Their offspring were also acclimated to cool and warm temperatures so that offspring environments matched or mismatched that of their mother and/or father. This allowed us to observe transgenerational effects when offspring and parental environments matched and compare them with effects when temperature conditions differed between generations. Because the mothers and fathers were from matched or mismatched environments, this provided us with an opportunity to assess the relative parental contribution of the parents to offspring thermal performance. Given the evidence supporting anticipatory effects from both mothers and fathers (Marshall, 2015; Shama *et al*., 2016), we hypothesized that parents additively contribute to transgenerational plasticity.

## Methods

These experiments were conducted in accordance with the guidelines of the Canadian Council on Animal Care. They have been approved by the Institutional Animal Care Committee of Trent University (Protocol # 24794) and the Ontario Ministry of Natural Resources and Forestry (OMNRF) Aquatic Animal Care Committee (Protocol # 136).

The strain of lake trout used in this experiment originated from Seneca Lake which is a glacial lake located in the Finger Lakes region in central New York state (42°41’ N, 76°54’ W) and has been kept in the OMNRF hatchery system since 1990 (OMNRF Fish Culture Stocks Catalogue 2005).

### Experimental design: Adult trout acclimation and breeding

Mature adult lake trout (age 8; 2.3 - 4.2 kg) were held at the OMNRF White Lake Fish Culture Station (Sharbot Lake, Ontario, Canada) where they were individually PIT tagged (Oregon RFID, Portland OR), divided into two groups (n = 8 and 9, mixed sex) and acclimated to two different temperatures (10 ± 0.5 °C and 17 ± 0.5 °C) beginning in July, 2015. The lower target temperature was based on lake trout temperature requirements for spawning and the elevated temperature was chosen to exceed their typical range but remaining within physiological limits (Casselman, 2008) with the aim of inducing a physiological stress response due to warming while attempting to avoid reproductive failure. Adults were housed in 1 × 1 × 6 m tanks that were covered with black tarpaulin to block out light. Temperatures were maintained by drawing water from above and below the thermocline in the hatchery’s water source (White Lake) and mixing it as it was fed into the tanks where the fish were held. After September, the temperature of each tank was allowed to follow the seasonal cooling of the lake.

Beginning in October, offspring were produced by dry-spawning anaesthetized fish (anaesthetic: 0.1 g/L MS-222; Aqua Life, Syndel Laboratories Ltd., B.C., Canada) where 140 mL of eggs were stripped from each female, divided evenly among 4 jars and fertilized by pipetting milt directly onto them. Families were produced by a full factorial mating cross using two males and two females from each of the two temperature treatments (8 fish in total) so that resultant offspring were from parents who had been acclimated to either the same or different temperatures prior to spawning. This resulted in four offspring families from each of the four parental treatment groups (W_♀_xW_♂_, W_♀_xC_♂_, C_♀_xW_♂_ and C_♀_xC_♂_, where W refers to a warm acclimated parent and C refers to a cold acclimated parent) for a total of 16 families. After fertilization, egg jars were kept cool and transported in coolers to the Codrington Fish Research Facility (Codrington, Ontario, Canada). Upon arrival, the eggs from each jar were placed in perforated steel boxes (9 × 9 × 7.5 cm, one family per box) which were kept in flow-through tanks receiving freshwater at ambient temperature (5-6 °C) and natural photoperiod under dim light.

### Experimental design: Offspring temperature acclimation

In March, when the fry reached the exogenous feeding stage, 14 individuals from each family were randomly selected, split into two groups of 7 and transferred into one of four larger (200 L) tanks. Each tank was separated into four sections to keep the families separate, however, due to space constraints two families were kept in each tank section where the offspring from the two families sharing a section were half-siblings by their father. The individuals would later be identified to family using microsatellite genotyping (Supplemental File 1). Two tanks received a cold/optimal temperature (11 °C) and the other two received a warm temperature (15 °C) so that each family had 7 representatives acclimated to each temperature. The lower acclimation temperature was selected based on the optimal growth temperature for lake trout (Edsall, 2002; Casselman, 2008), and the warm temperature represents the potential warming in the Great Lakes region due to climate change by the end of the century (Hayhoe *et al*., 2010).

After transferring the fry to the larger tanks, we changed the water temperature at a rate of 1 °C per day until the target temperatures (11 and 15 °C) were reached, and the fish were acclimated for 3-4 weeks before the experiments began. The fish were fed 5-6 times a day at 2-3% their mass, however, fish were fasted for at least 12 hours prior to experimentation so that the physiological effects from recent feeding did not influence experimental results (Millidine *et al*., 2009).

### Respirometry set up

To test for potential transgenerational effects of the parental environment on offspring physiology, we measured and compared the metabolic rate of offspring of parents acclimated to matched or mismatched temperature conditions. To do so, we first measured the metabolic rate of offspring as the rate of oxygen consumption (MO_2_), using closed respirometry during an acute temperature increase. From this dataset, we determined each individual offspring’s standard rate of oxygen consumption (MO_2_) and peak MO_2_. The standard MO_2_ was recorded as the MO_2_ at the fish’s acclimation temperature before temperature began to rise with the acute temperature challenge (Chabot *et al*., 2016), and the peak MO_2_ was recorded as the highest MO_2_ achieved during the trial. To determine the upper thermal tolerance in the offspring, we measured the critical thermal maximum (CTM) which is the highest temperature that can be tolerated by the fish. This was recorded as the temperature at which the fish was no longer able to maintain equilibrium as temperature increased.

Respirometers consisted of custom-built glass cylinders (8 cm diameter × 4.5 cm height) sealed at one end and fitted with an acrylic lid. Each lid had an inlet and outlet valve to allow water to flow through the chambers using a submersible pump that circulated water through the respirometers at 4.5 L min^−1^. The valves were situated on either side of a fitting that held a dissolved oxygen probe (model DO-BTA, Vernier Software and Technology, OR, USA) in place. The respirometers were contained in clear plastic tubs (two respirometers per tub) atop two side-by-side stir plates so that each respirometer was positioned over a stir plate. A magnetic stir bar in each respirometer was set to spin at approximately 60 RPM to keep water circulating in the chamber and a perforated stainless-steel grid separated the fish from the stir bar. The containers received aerated freshwater from a source tank that was temperature controlled using three 500W titanium heaters (model TH-0500, Finnex, IL, USA) with digital temperature controllers (model HC 810M, Finnex). The plastic tubs were covered in a sheet of thin, black plastic to minimize visual disturbance to the fish.

### Respirometry protocol and determining critical thermal maximum

The night before the experiment, eight fish were individually transferred into separate respirometers where they received a continuous flow of fresh water maintained at their acclimation temperature and were left to adjust to the experimental apparatus overnight. Standard MO_2_ was measured the following morning. MO_2_ measurements were collected by manually switching off the pumps that circulated water through the respirometers and closing the input and output valves to create a closed system. The stir bar kept water moving past the oxygen probe which was connected to a Lab Pro (Vernier Software and Technology) interfaced with LoggerPro software (version 3.8.6; Vernier Software and Technology) so that the reduction in oxygen concentration could be recorded. Measurement of MO_2_ began after a 30 second wait period, then the drop in O_2_ was recorded for 10 minutes, after which the valves were opened to allow fresh water to flush the chamber until the next oxygen consumption measurement was made (approximately 30 minutes).

After measurement of standard MO_2_, fish were subjected to an acute temperature challenge of +2 °C per hour by raising the temperature of the water in the source tank that fed the tubs housing the respirometers. We choose this rate to be consistent with previous studies that measured metabolic rate via oxygen consumption in related species (Penney *et al*., 2014). We measured MO_2_, for 10 minutes, at every 1 °C increase until the fish lost equilibrium which was observed when the fish could no longer maintain an upright position in the respirometer chamber, and this was recorded as the CTM for that fish. At this point, the focal fish was quickly removed from the chamber and euthanized in a bath of 0.3 g/L of tricaine methanesulfonate (MS-222; Aqua Life, Syndel Laboratories Ltd., B.C., Canada). The focal fish was blotted dry on a paper towel so that mass (measured to the nearest 0.1 g using a digital balance scale) and fork length (measured to the nearest 1 mm using digital calipers) could be measured, and a caudal fin tissue sample (~0.25cm^2^) was preserved in 95% ethanol for subsequent genotyping to identify offspring individuals to their respective family (see supplementary material).

The oxygen saturation of the water in the source tank and respirometers was continuously monitored throughout each trial. The measurement period was shortened if oxygen concentration in the respirometers began to approach the critical limit of 3.5 mg O_2_ L^−1^ (Doudoroff & Shumway, 1970; Cook *et al*., 2018). Also, if the oxygen saturation levels in the source began to drop due to higher temperatures, then oxygen was supplemented to the source tank water with a tank and diffuser.

### Calculations and statistical analysis

Reduction in oxygen concentration was recorded as mg O_2_ L^−1^ min^−1^, and the rate of oxygen consumption (MO_2_) was calculated using the following formula,

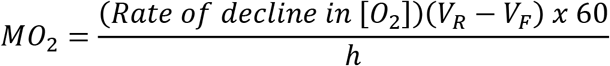

where *(Rate of decline in [O_2_])* is the decline in water oxygen concentration during the 10-minute measurement period, *V*_*R*_ is the volume (L) of the respirometers, *V*_*F*_ is the volume of the fish (L) and *h* is the time in hours.

Condition factor was calculated as,

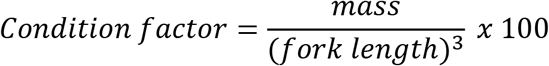

To explore factors that contributed to variation in body mass and condition factor we used using JMP 13 (v. 18.1). Statistical analyses of the MO_2_ during the temperature challenge, and the standard and peak MO_2_ were conducted using R (v. 3.5.2) with the ‘MuMIn’ (Barton, 2019), ‘lme4’ (Bates *et al*., 2015), and ‘mgcv’ (Wood, 2011) packages. The level of significance was set to 0.05 in all analyses, and all model assumptions (linearity, homogeneity of variance, sample independence, and residual normality) were confirmed visually, and by use of Shapiro-Wilk W, Levene’s and the Brown-Forsythe tests. In some cases, our response variable appeared non-normally distributed (according to Shapiro-Wilk W tests), however, we still opted for parametric tests as our selected analytical approaches are not highly sensitive to non-normality (Glass *et al*., 1972; Harwell *et al*., 1992; Lix *et al*., 1996; Bodden *et al*., 2017; Senduran *et al*., 2018) and depend more on homogeneity of variance instead.

To test for an effect of maternal, paternal and offspring acclimation temperatures on offspring condition factor and mass we used two separate general linear mixed effects models (GLMM) in JMP, with mass and condition factor as Gaussian-distributed response variables. These models both included offspring acclimation temperature (cold or warm) and parent acclimation temperature as fixed effect predictors, where parents were treated as a single explanatory variable with mother and father acclimation temperature combined and represented as one of four fixed effects: C_♀_xC_♂_, C_♀_xW_♂_, W_♀_xC_♂_ or W_♀_xW_♂_ (C = cold acclimation and W = warm acclimation). To test for whether parental acclimation temperature yielded differential effects on mass and condition of offspring reared in cold or warm water, an interaction between offspring and parental treatment group was also included as a fixed effect predictor. Degree days was included as a random intercept to control for effects of age on mass and condition, since the experiment lasted approximately five weeks and most of the cold acclimated offspring were tested in the first half of the experimental period. Here, degree days were calculated for each fish as the cumulative temperature experienced above 0 °C (Chezik *et al*., 2013; Cook *et al*., 2018) until the day of the experiment. Finally, offspring identity (ID) and parental IDs (*IDM* and *IDF*) were also included as random intercepts to account for statistical non-independence between offspring that were sired or dammed from the same parents, and between repeated measures drawn from the same individual.

To test the effect of maternal, paternal and offspring acclimation temperature on the metabolic (MO_2_) response of the offspring to an acute temperature challenge, we again used a GLMM, however, here, our model was constructed using the ‘nlme’ package in R (Pinheiro et al, 2019) to permit correction for temporal autocorrelation. In this model, MO_2_ was used as a Gaussian-distributed response variable, with acute challenge temperature (*T*_*a*_; continuous variable), offspring acclimation temperature (*T*_*O*_; cold and warm), and the acclimation temperature of the mothers (*T*_*M*_; cold and warm) and the fathers (*T*_*F*_; cold and warm) used as fixed effect predictors, along with all possible interactions between these terms. Additionally, *Mass* (fixed term) was also included in the model as a continuous predictor since the warm acclimated offspring grew heavier than cold acclimated ones, and because metabolic rate scales with mass. Similar to our previous models, both mother and father ID (*ID*_*M*_ and *ID*_*F*_) were included as random intercepts to account for the relatedness among the offspring, and offspring ID was also included as a random intercept to control for statistical non-independence between measurements drawn from the same individual.

Because the relationship between MO_2_ and acute temperature challenge was curvilinear and could not be predicted by a simple polynomial function (i.e. with relatively low degree), we first modeled the relationship between MO_2_ and acute challenge temperature alone using a cubic regression spline in a general additive model (GAM), using three knots to appropriately capture the shape of the relationship while avoiding over-fitting. Predicted MO_2_ at each challenge temperature was extracted for each offspring and used in place of acute challenge temperature within our GLMM to account for the variation in the response variable (observed MO_2_) due to acute challenge temperature. This approach permitted us to: 1) remove the complex, curvilinear relationship between MO_2_ and acute challenge temperature, 2) test whether the remaining variation in MO_2_ (i.e., that not explained by acute challenge temperature) can be explained by the other terms in the GLMM, and 3) include multi-level interactions between a previously non-linear predictor (acute challenge temperature) and additional factorial predictors, which cannot be accomplished simply with current additive models. Finally, to account for heterogeneity of variance in MO_2_ across acute challenge temperature (and detected across predicted MO_2_), and to correct for autocorrelation between measurements drawn at adjacent time-points, we weighted our model by acute challenge temperature, and included a type I autoregressive correlation structure, with an estimated ρ of 0.397.

An effect of parental acclimation temperature on the standard MO_2_, peak MO_2_ and CTM of offspring were analyzed using three independent linear mixed models in R, each with the ‘lme4’ package (Bates et al, 2015). Here, we first sought to include mother acclimation temperature (*T*_*M*_), father acclimation temperature (*T*_*F*_), and offspring acclimation temperature (*T*_*O*_) as fixed effect predictors, with all possible interactions between each of these factors, along with offspring *mass* as a covariate. Unfortunately, however, our total number of observations per experimental group (n̄ = 21.125 ± s.d. = 2.642; total n = 157) were too few to support a robust test regarding the individual effects of each predictor (π = 0.448; for expected relationships with weak explanatory capacity; Cohen’s f^2^ ≅ 0.05; as tested using the ‘pwr’ package in r; Champely et al, 2018). We therefore used an Akaike Information Criterion (AIC) approach to identify which of the models best explained the variation in the data while avoiding over-parameterization, using the previously described global model, including mother ID (*ID*_*M*_) and father ID (*ID*_*F*_) as random intercepts to account for the relatedness among the offspring. The best models were considered as those with a ΔAIC ≤ 2 as recommended by Burnham and Anderson (2002) where the ΔAIC was calculated as the difference in the AIC value of a given model versus the top model (i.e. the model with the lowest AIC value). We also calculated the evidence ratio (ER) and Akaike weight (Wi) of each model iteration. The ER is the likelihood that the top model is the best supporting model compared to another model, and the Wi is the weight (or proportion) of evidence that a given model is best explains the variation in the data (Burnham & Anderson, 2002). We used these metrics to compare the best models and to observe common parameters among these models.

If cold-adapted, stenothermal ectotherms can better cope with environmental warming through transgenerational plasticity, we predict that the MO_2_ of offspring from warm acclimated parents would be higher at all challenge temperatures compared to those from cold acclimated parents (Fig. 1). Any interactions that occur between the variables will be observed as a crossing of the lines. Also, warm acclimated offspring would tolerate higher temperatures, as indicated by a higher CTM, and attain a higher standard and peak MO_2_ (Fig. 1). Statistically this would be observed as a significant effect of parental temperature (*T*_*M*_ or *T*_*F*_). If the hypothesis that the parents additively contribute to the transgenerational effect is correct, then both parental terms (*T*_*M*_ and *T*_*F*_) will have a significant effect on the response variables.

**Figure 1:**
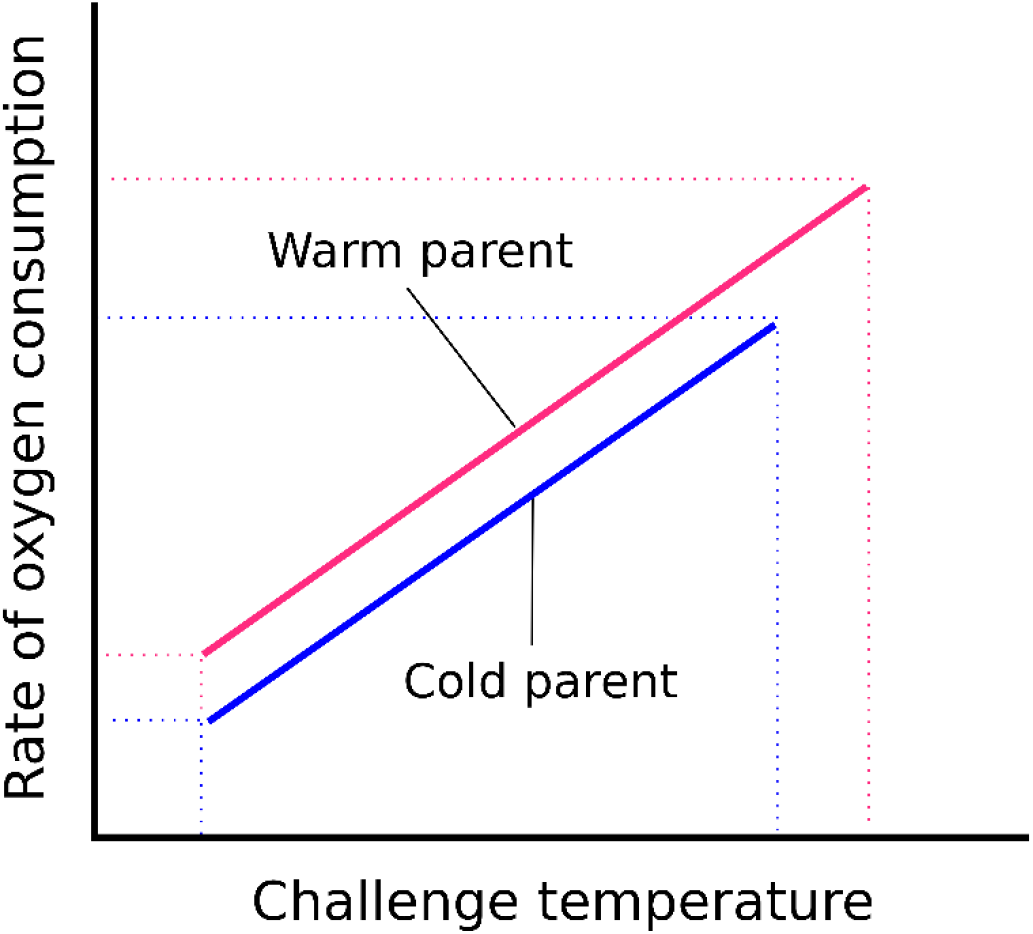
The predicted effect parent acclimation temperature on the rate of oxygen consumption of their offspring. The effect could be driven by either the maternal or paternal acclimation temperatures. Interactions would be observed as a crossing of the lines.

## Results

### Mass and condition factor

On average, warm acclimated offspring were nearly twice as heavy compared with cold acclimated offspring (least squares means: 4.28 ± 0.14 g versus 2.76 ± 0.13 g; GLMM: F_1,21.28_ = 93.58, p < 0.001; Table 1). Offspring mass did not differ among parental groupings (C_♀_xC_♂_, C_♀_xW_♂_, W_♀_xC_♂_, W_♀_xW_♂_) (GLMM: F_3,3.63_ = 1.08, p = 0.45), but there was an interaction between offspring acclimation and parental group (GLMM: F_3,46.53_ = 4.23, p = 0.01).

**Table 1:** The mass and condition factor of 11 °C and 15 °C acclimated lake trout offspring. Parental groups are represented as the maternal environment crossed with the paternal environment: C_♀_xC_♂_, C_♀_xW_♂_, W_♀_xC_♂_ and W_♀_xW_♂_ where C = cold and W = warm. Values are least squares means ± SEM. Statistical significance (p < 0.05) between offspring acclimation temperature is indicated by an asterisk.

| Parental group | 11 °C Acclimated offspring |  |  |  | 15 °C Acclimated offspring |  |  |  |
| --- | --- | --- | --- | --- | --- | --- | --- | --- |
|  | C♀x C♂ | C♀x W♂ | W♀x C♂ | W♀x W♂ | C♀x C♂ | C♀x W♂ | W♀x C♂ | W♀x W♂ |
| Mass (g) | 2.93<br>($\pm 0.26$) | 2.69<br>($\pm 0.23$) | 3.26<br>($\pm 0.22$) | 2.14<br>($\pm 0.26$) | 4.16*<br>($\pm 0.25$) | 4.61*<br>($\pm 0.28$) | 4.17*<br>($\pm 0.25$) | 4.19*<br>($\pm 0.27$) |
| Condition factor | 0.91<br>($\pm 0.01$) | 0.88<br>($\pm 0.01$) | 0.91<br>($\pm 0.01$) | 0.86<br>($\pm 0.01$) | 0.93<br>($\pm 0.01$) | 0.94<br>($\pm 0.01$) | 0.94<br>($\pm 0.01$) | 0.92<br>($\pm 0.01$) |

Warm acclimated offspring had significantly higher body condition than cold acclimated offspring with means of 0.93 ± 0.01 vs. 0.89 ± 0.01 (Table 1), respectively (GLMM: F_1,20.47_ = 38.67, p < 0.001). Offspring condition factor was not affected by parent acclimation temperatures (GLMM: F_3,3.23_ = 1.83, p = 0.31) nor by the interaction between offspring acclimation and parental group (GLMM: F_3,2.07_, p = 0.12).

### Metabolic response of offspring to an acute temperature challenge

There was an increase in offspring MO_2_ with increasing body mass (*Mass*: t = 10.66, p < 0.001, Table 2). Offspring MO_2_ also increased with challenge temperature (GAMM: *T*_*a*_: t = 17.58, p < 0.001). Offspring acclimation temperature (*T*_*O*_) had a significant effect on MO_2_ with warm acclimated offspring having a higher MO_2_ (*T*_*O*_: t = −3.40, p < 0.001, Table 2). Neither maternal nor paternal acclimation temperature in isolation was strong enough to influence offspring MO_2_ (*T*_*M*_: t = 2.12, p = 0.068; *T*_*F*_: t = 1.22, p = 0.222; Table 2).

**Table 2:**
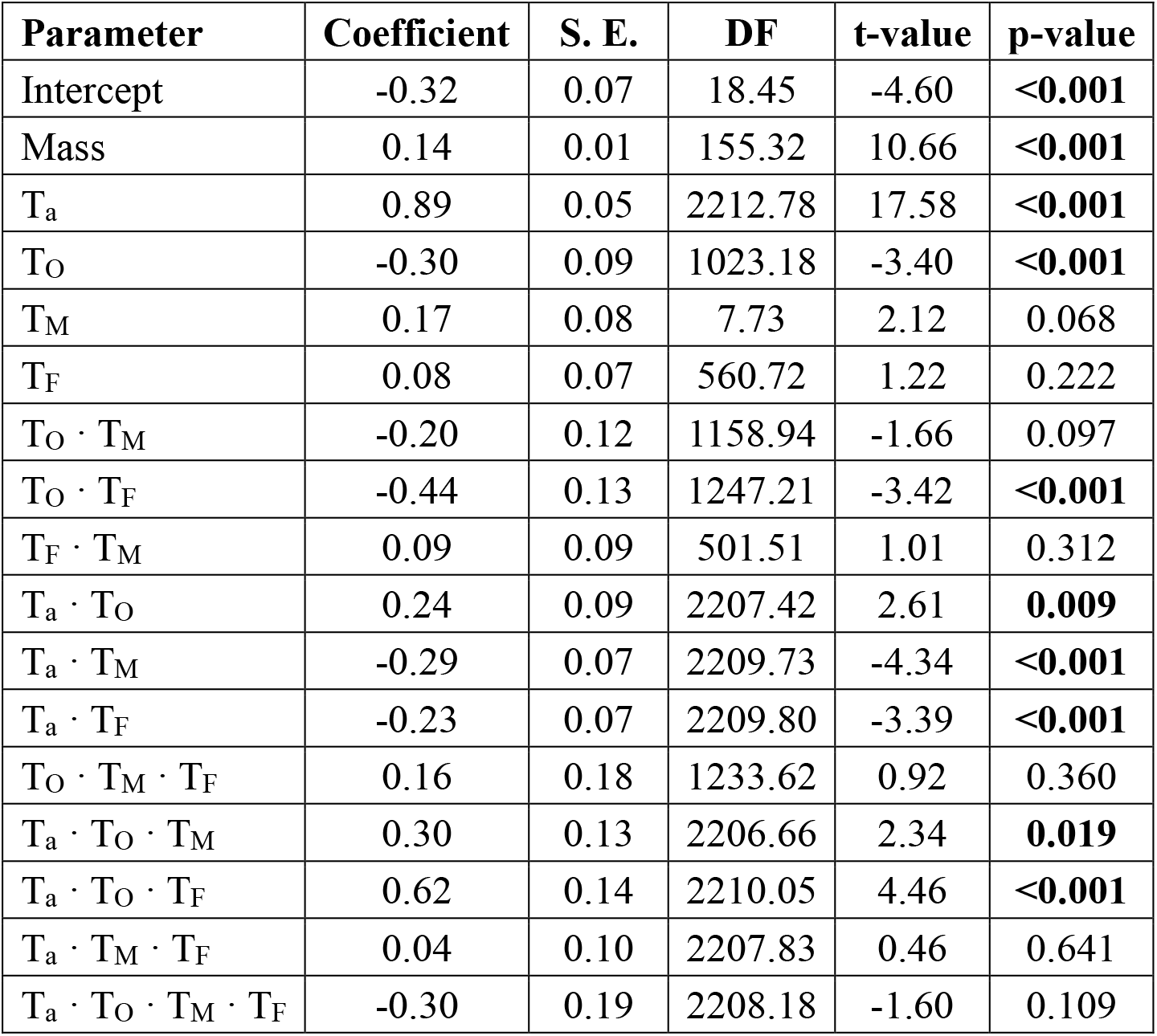
Summary of the GLMM results to test for a transgenerational effect of acclimation temperature on lake trout offspring MO_2_ during an acute temperature challenge. Offspring were from parents acclimated to either a cold or warm temperature. The offspring were also acclimated to cold or warm temperature. *T*_*F*_, *T*_*M*_, and *T*_*O*_ are the father, mother and offspring acclimation temperatures, respectively, and the acute temperature challenge is represented by *T*_*a*_. Significant effects (p<0.05) are highlighted with bold text.

| Parameter | Coefficient | S. E. | DF | t-value | p-value |
| --- | --- | --- | --- | --- | --- |
| Intercept | -0.32 | 0.07 | 18.45 | -4.60 | **<0.001** |
| Mass | 0.14 | 0.01 | 155.32 | 10.66 | **<0.001** |
| $T_a$ | 0.89 | 0.05 | 2212.78 | 17.58 | **<0.001** |
| $T_O$ | -0.30 | 0.09 | 1023.18 | -3.40 | **<0.001** |
| $T_M$ | 0.17 | 0.08 | 7.73 | 2.12 | 0.068 |
| $T_F$ | 0.08 | 0.07 | 560.72 | 1.22 | 0.222 |
| $T_O \cdot T_M$ | -0.20 | 0.12 | 1158.94 | -1.66 | 0.097 |
| $T_O \cdot T_F$ | -0.44 | 0.13 | 1247.21 | -3.42 | **<0.001** |
| $T_F \cdot T_M$ | 0.09 | 0.09 | 501.51 | 1.01 | 0.312 |
| $T_a \cdot T_O$ | 0.24 | 0.09 | 2207.42 | 2.61 | **0.009** |
| $T_a \cdot T_M$ | -0.29 | 0.07 | 2209.73 | -4.34 | **<0.001** |
| $T_a \cdot T_F$ | -0.23 | 0.07 | 2209.80 | -3.39 | **<0.001** |
| $T_O \cdot T_M \cdot T_F$ | 0.16 | 0.18 | 1233.62 | 0.92 | 0.360 |
| $T_a \cdot T_O \cdot T_M$ | 0.30 | 0.13 | 2206.66 | 2.34 | **0.019** |
| $T_a \cdot T_O \cdot T_F$ | 0.62 | 0.14 | 2210.05 | 4.46 | **<0.001** |
| $T_a \cdot T_M \cdot T_F$ | 0.04 | 0.10 | 2207.83 | 0.46 | 0.641 |
| $T_a \cdot T_O \cdot T_M \cdot T_F$ | -0.30 | 0.19 | 2208.18 | -1.60 | 0.109 |

While the interaction between offspring and maternal acclimation temperature was not significant (*T*_*O*_ · *T*_*M*_: t = −1.66, p = 0.097), the interaction between offspring and paternal acclimation temperature did influence MO_2_ (*T*_*O*_ · *T*_*F*_: t = −3.42, p < 0.001). There was no significant interaction between mother and father acclimation temperature on the offspring’s metabolic response (*T*_*M*_ · *T*_*F*_: t = 1.01, p = 0.312). Significant two-way interactions occurred between *T*_*a*_ and *T*_*O*_ (t = 2.61, p = 0.009) demonstrating that some remaining variation in MO_2_ that was not explained by challenge temperature could be explained by offspring acclimation temperature; more specifically, offspring reared at warm temperatures appeared to respond differently to the thermal challenges than did cold acclimated offspring. The acclimation temperature of the parents interacted significantly with the acute temperature challenge to determine the offspring’s MO_2_ (*T*_*a*_ · *T*_*M*_: t = −4.34, p < 0.001; 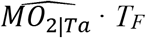: t = −3.39, p < 0.001; Table 2). The metabolic response of the offspring also depended on the complex interaction between offspring acclimation temperature, parental (maternal or paternal) acclimation temperature and challenge temperature (*T*_*a*_ · *T*_*O*_ · *T*_*M*_, t = 2.34, p = 0.019; *T*_*a*_ · *T*_*O*_ · *T*_*F*_, t = 4.46, p < 0.001; Table 1). No other main effect interactions were significant (Table 2).

To visually explore maternal and paternal influences on offspring MO_2_, we plotted the mass-specific MO_2_ for offspring from different parental combinations against the acute temperature challenge (Fig. 2). We did not perform a statistical analysis on the mass-specific values because the GLMM (previously described) tested MO_2_ while accounting for mass in the model. Qualitatively, the cold acclimated offspring from warm acclimated parents (W_♀_xW_♂_) had a higher metabolic rate at the beginning of the temperature challenge compared with the other parental acclimation groups (C_♀_xC_♂_, C_♀_xW_♂_, W_♀_xC_♂_; Fig. 2A) indicating that an environmental mismatch between generations can influence offspring metabolic response. This effect did not carry over to the warm acclimated offspring (Fig. 2B) as MO_2_ was comparable among the parental acclimation groups.

**Figure 2:**
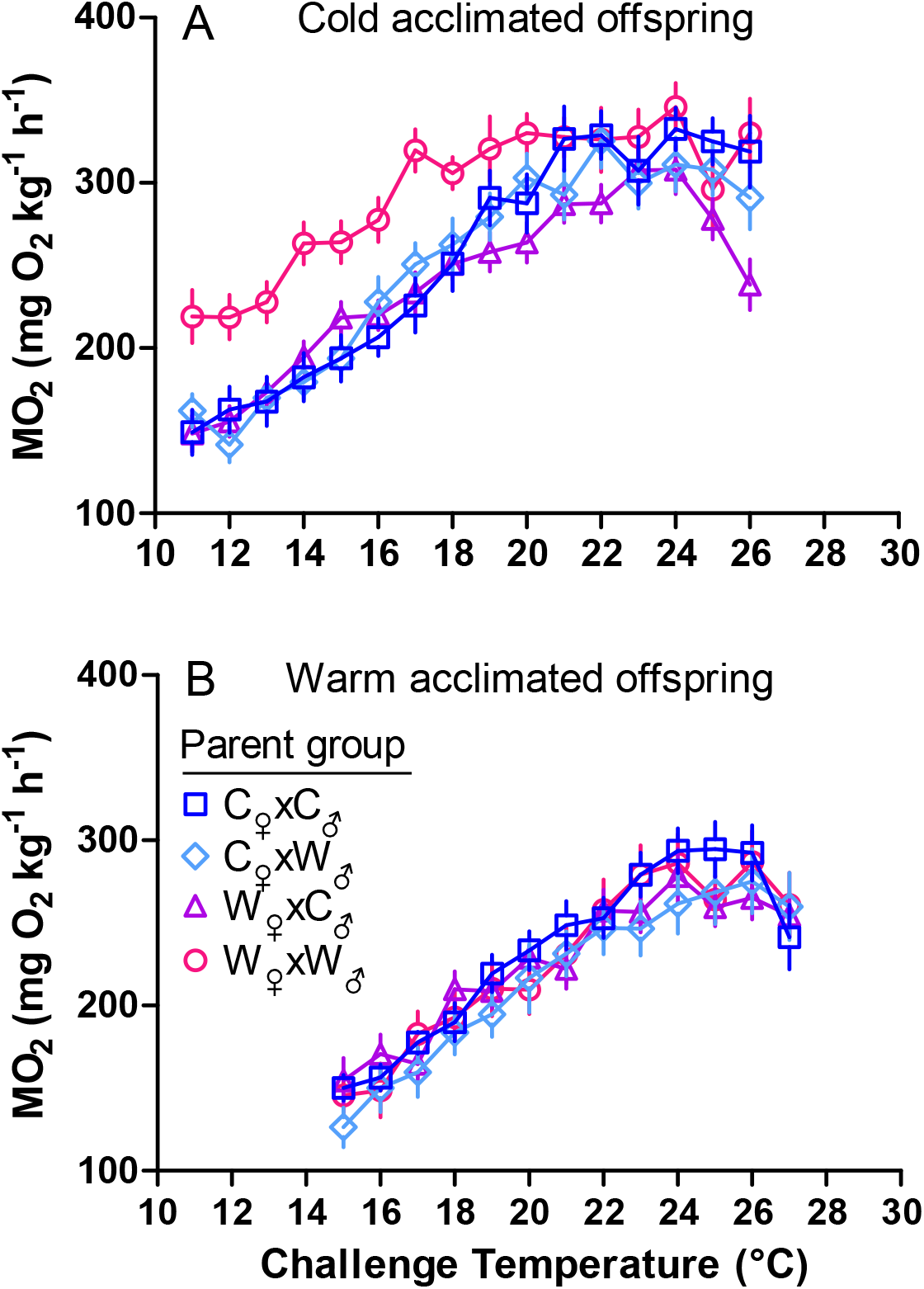
The change in the rate of oxygen consumption (MO_2_) of A) cold and B) warm acclimated lake trout offspring given an acute temperature challenge of +2°C·h^−1^, showing mass-specific means ± SEM. Parental groups are represented as the maternal environment crossed with the paternal environment: C_♀_xC_♂_, C_♀_xW_♂_, W_♀_xC_♂_ and W_♀_xW_♂_ where C = cold and W = warm.

To visually isolate the effects of maternal acclimation temperature on offspring’s thermal response, we plotted the mass-adjusted MO_2_ (Fig. 3) estimated from the GLMM to show the interaction between challenge temperature and the acclimation temperature of the mothers (*T*_*a*_ · *T*_*M*_; Table 2). For both cold and warm acclimated offspring (Fig. 3, both panels) the difference in the slope of the reaction norms illustrates the significant interaction between challenge temperature and acclimation temperatures of the offspring and mothers (*T*_*a*_ · *T*_*O*_ · *T*_*M*_; p = 0.019; Table 2). Focusing on the cold acclimated offspring (Fig. 3, left), at cooler challenge temperatures, the MO_2_ of offspring from warm acclimated mothers was elevated compared to offspring from cold acclimated mothers. For warm acclimated offspring at challenge temperatures below approximately 19 °C, individuals from warm acclimated mothers had a higher MO_2_ compared to those from cold acclimated mothers (Fig. 3A). This general trend occurred to a lesser extent in the warm acclimated offspring (Fig. 3B) with the offspring’s MO_2_ reaction norms overlapping for warm and cold acclimated mothers.

**Figure 3:**
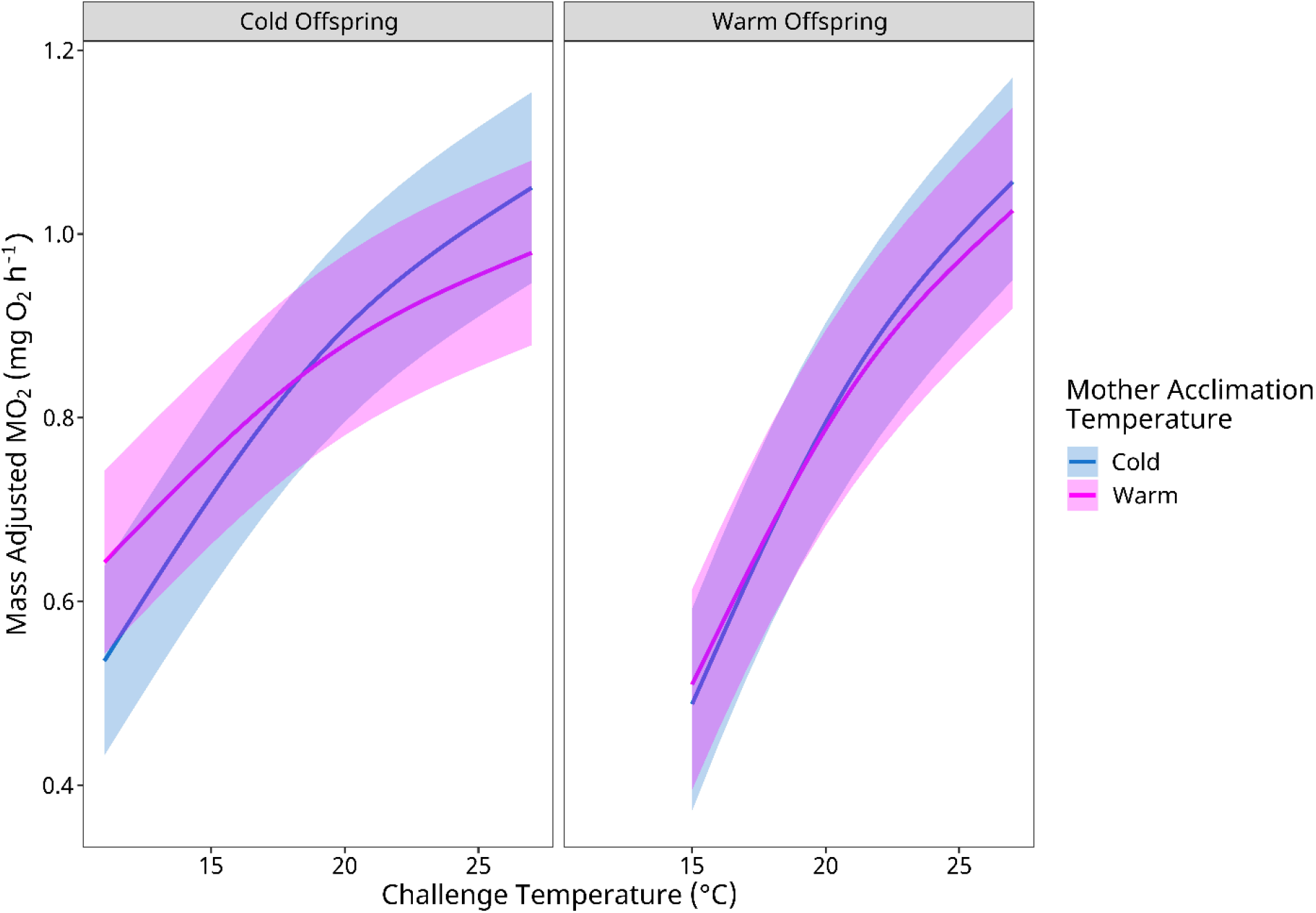
The influence of maternal acclimation temperature on the change in the rate of oxygen consumption (MO_2_) of cold and warm acclimated lake trout offspring given an acute temperature challenge of +2°C·h^−1^. Values are means estimated from the GLMM with 95% confidence intervals (refer to Methods).

To visually explore the paternal effect on offspring MO_2_ we plotted the mass-adjusted MO_2_, estimated from the GLMM (Fig. 4). This illustrates the significant interaction between challenge temperature and the acclimation temperatures of the offspring and fathers (*T*_*a*_ · *T*_*O*_ · *T*_*F*_; p < 0.001; Table 2). For cold acclimated offspring, the MO_2_ reaction norm of those from warm acclimated fathers was above that of those from cold acclimated fathers with the lines crossing at approximately 14 °C (Fig. 4A). A reverse trend occurred in the warm acclimated offspring as individuals from warm acclimated fathers had a lower MO_2_ at cooler challenge temperatures compared to those from cold acclimated fathers (Fig. 4B).

**Figure 4:**
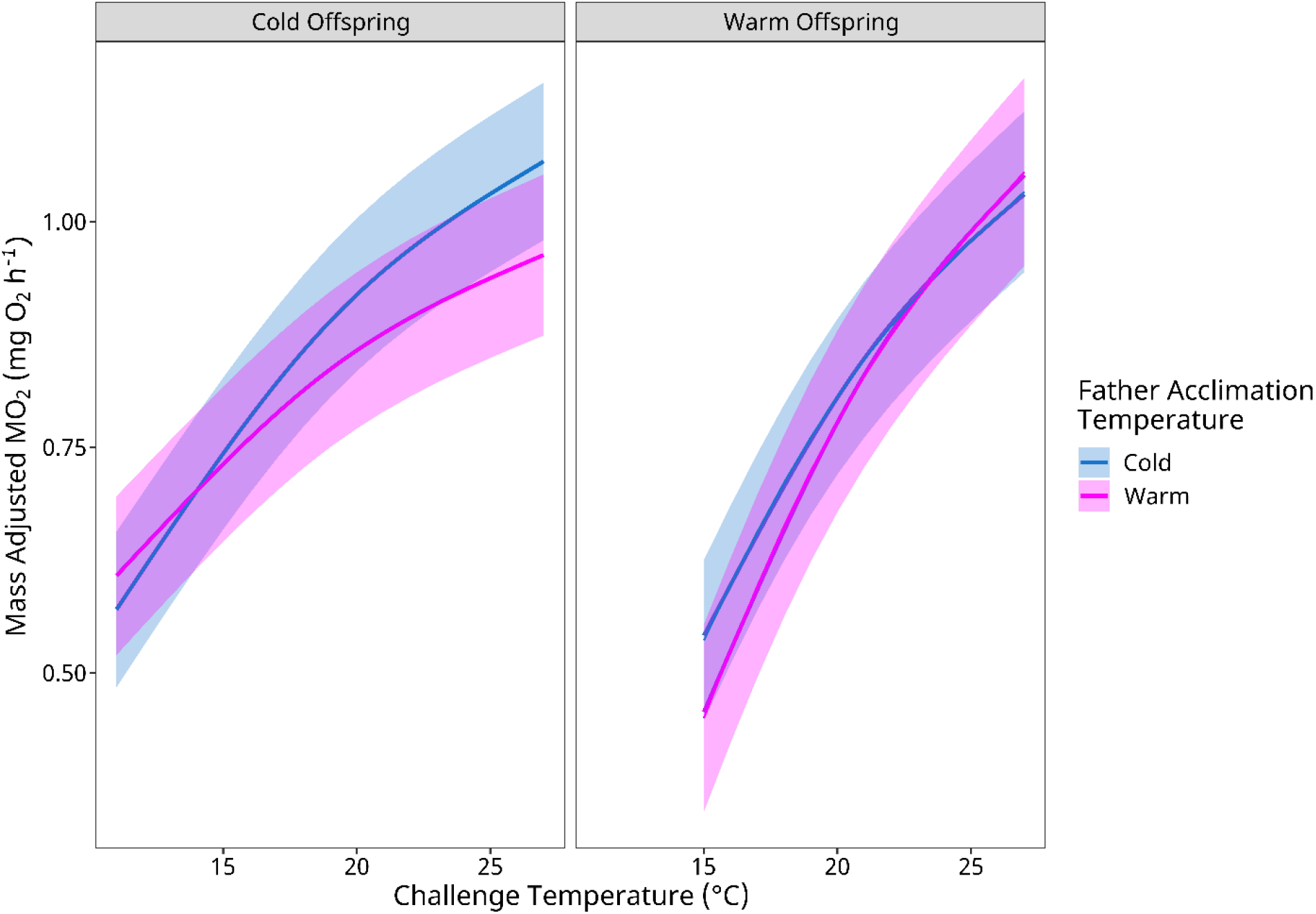
The influence of paternal acclimation temperature on the change in the rate of oxygen consumption (MO_2_) of cold and warm acclimated lake trout offspring given an acute temperature challenge of +2°C·h^−1^. Values are means estimated from the GLMM with 95% confidence intervals (refer to Methods).

### Standard and peak metabolic rate

Analysis of standard MO_2_ with AIC revealed six models (ΔAIC ≤ 2) that best predicted the trends in the data with *Mass* appearing in each of these models. The first model contained *Mass* as the only fixed variable, but this model was only 1.23 times more likely (evidence ratio, ER) than model 2, which included maternal (*T*_*M*_) and paternal (*T*_*F*_) acclimation temperature with their interaction, to best explain the variation in the data (Table 3). Interestingly, offspring acclimation temperature (*T*_*O*_) appeared only twice among the six models. Maternal (*T*_*M*_) and paternal (*T*_*F*_) acclimation temperature appeared most frequently among the six models, with an interaction between these terms appearing in two of these. An interaction between offspring and paternal acclimation temperature (*T*_*O*_ · *T*_*F*_) appeared in only one of the models (Table 3).

**Table 3:**
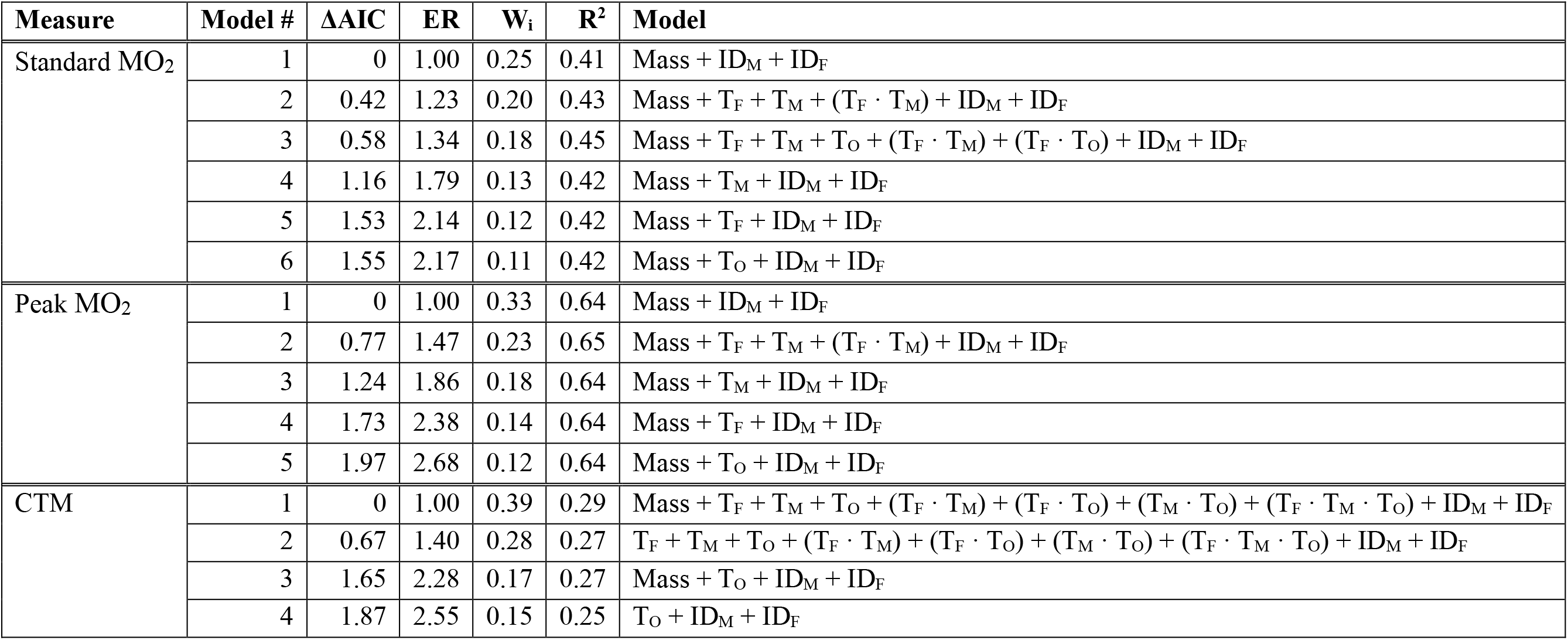
Summary of the top models determined with AIC to explain variation in standard MO_2_, peak MO_2_ and critical thermal maximum (CTM) with transgenerational acclimation of lake trout offspring. Offspring were from parents acclimated to either a cold or warm temperature and were also acclimated to cold or warm temperature. *T*_*F*_, *T*_*M*_, and *T*_*O*_ are the father, mother and offspring acclimation temperatures, respectively, and IDM and IDF are the mother and father individual identification (treated as random effects).

| Measure | Model # | $\Delta\text{AIC}$ | ER | $W_i$ | $R^2$ | Model |
| --- | --- | --- | --- | --- | --- | --- |
| Standard $\text{MO}_2$ | 1 | 0 | 1.00 | 0.25 | 0.41 | Mass + $\text{ID}_M$ + $\text{ID}_F$ |
| | 2 | 0.42 | 1.23 | 0.20 | 0.43 | Mass + $T_F$ + $T_M$ + ($T_F \cdot T_M$) + $\text{ID}_M$ + $\text{ID}_F$ |
| | 3 | 0.58 | 1.34 | 0.18 | 0.45 | Mass + $T_F$ + $T_M$ + $T_O$ + ($T_F \cdot T_M$) + ($T_F \cdot T_O$) + $\text{ID}_M$ + $\text{ID}_F$ |
| | 4 | 1.16 | 1.79 | 0.13 | 0.42 | Mass + $T_M$ + $\text{ID}_M$ + $\text{ID}_F$ |
| | 5 | 1.53 | 2.14 | 0.12 | 0.42 | Mass + $T_F$ + $\text{ID}_M$ + $\text{ID}_F$ |
| | 6 | 1.55 | 2.17 | 0.11 | 0.42 | Mass + $T_O$ + $\text{ID}_M$ + $\text{ID}_F$ |
| Peak $\text{MO}_2$ | 1 | 0 | 1.00 | 0.33 | 0.64 | Mass + $\text{ID}_M$ + $\text{ID}_F$ |
| | 2 | 0.77 | 1.47 | 0.23 | 0.65 | Mass + $T_F$ + $T_M$ + ($T_F \cdot T_M$) + $\text{ID}_M$ + $\text{ID}_F$ |
| | 3 | 1.24 | 1.86 | 0.18 | 0.64 | Mass + $T_M$ + $\text{ID}_M$ + $\text{ID}_F$ |
| | 4 | 1.73 | 2.38 | 0.14 | 0.64 | Mass + $T_F$ + $\text{ID}_M$ + $\text{ID}_F$ |
| | 5 | 1.97 | 2.68 | 0.12 | 0.64 | Mass + $T_O$ + $\text{ID}_M$ + $\text{ID}_F$ |
| CTM | 1 | 0 | 1.00 | 0.39 | 0.29 | Mass + $T_F$ + $T_M$ + $T_O$ + ($T_F \cdot T_M$) + ($T_F \cdot T_O$) + ($T_M \cdot T_O$) + ($T_F \cdot T_M \cdot T_O$) + $\text{ID}_M$ + $\text{ID}_F$ |
| | 2 | 0.67 | 1.40 | 0.28 | 0.27 | $T_F$ + $T_M$ + $T_O$ + ($T_F \cdot T_M$) + ($T_F \cdot T_O$) + ($T_M \cdot T_O$) + ($T_F \cdot T_M \cdot T_O$) + $\text{ID}_M$ + $\text{ID}_F$ |
| | 3 | 1.65 | 2.28 | 0.17 | 0.27 | Mass + $T_O$ + $\text{ID}_M$ + $\text{ID}_F$ |
| | 4 | 1.87 | 2.55 | 0.15 | 0.25 | $T_O$ + $\text{ID}_M$ + $\text{ID}_F$ |

Altogether this suggests that maternal and paternal environments, individually and combined, can act on the response of the offspring’s standard metabolic rate to temperature acclimation.

We plotted the standard MO_2_ (mass-specific values, no statistical analysis) to visually explore trends within and between the offspring and parental acclimation groups (Fig. 5A). The cold acclimated offspring from warm acclimated parents (open boxes, W_♀_xW_♂_, Fig. 5) had the highest mean standard MO_2_ (219.2 ± 15.97 mg O_2_ kg^−1^ h^−1^), while the standard MO_2_ of the other three groups ranged between 148.3 ± 11.08 and 162.1 ± 10.51 mg O_2_ kg^−1^ h^−1^ (Fig. 5A). There was no observable trend for warm acclimated offspring (shaded boxes, Fig. 5A) as standard MO_2_, irrespective of parental acclimation temperature, ranged between 130.4 ± 12.56 and 154.7 ± 13.52 mg O_2_ kg^−1^ h^−1^. When comparing offspring within parental acclimation temperatures (open vs. shaded boxes; Fig. 5A), the mean standard MO_2_ of cold acclimated offspring from warm acclimated parents (shaded boxes, W_♀_xW_♂_; Fig. 5A) was higher compared to cold acclimated offspring (open boxes, W_♀_xW_♂_; Fig. 5A).

**Figure 5:**
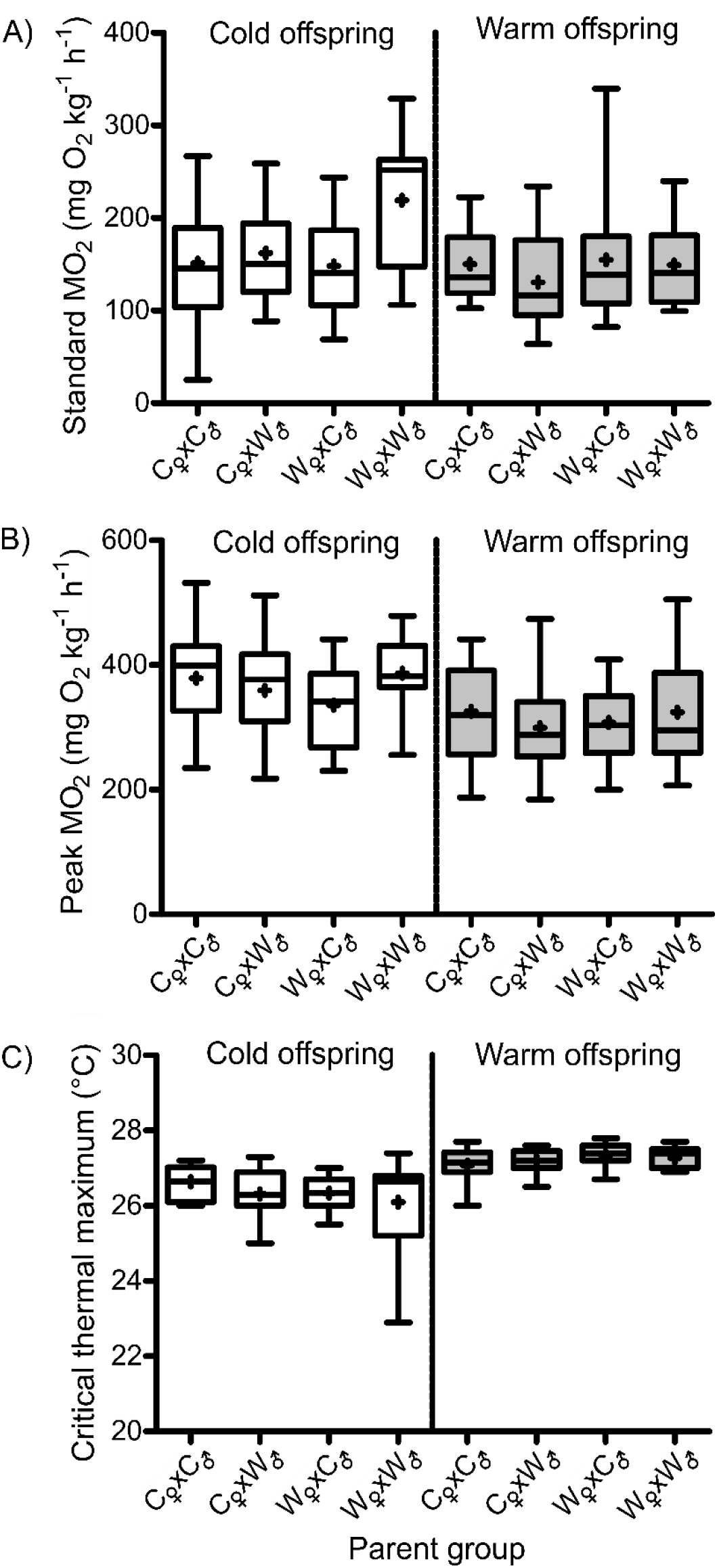
The A) standard rate of oxygen consumption (MO_2_), B) peak MO_2_ and C) critical thermal maximum (CTM) of lake trout offspring acclimated to a cold (open) or warm (shaded) temperature. Parental groups are represented as the maternal environment crossed with the paternal environment: C_♀_xC_♂_, C_♀_xW_♂_, W_♀_xC_♂_ and W_♀_xW_♂_ where C = cold and W = warm. The plot shows the 25^th^ and 75^th^ quartiles with the median, the mean is represented as ‘+’, and the upper and lower tails are the minimum and maximum values.

We also used an information theoretic approach to explore factors contributing to variation in peak MO_2_. Of the five models that best explained the variation in peak MO_2_, *mass* appeared in each model with the top model (ER = 1, W_i_ = 0.33) containing *mass* as the only fixed parameter (Table 3). Maternal (*T*_*M*_) and paternal (*T*_*F*_) acclimation temperature, and the interaction term between the two, occurred in the second-best model which had a 23% (W_i_) chance of being the top model (Table 3). The other three of the five models contained only one fixed parameter: either *T*_*O*_, *T*_*M*_, or *T*_*F*_ (Table 3).

Peak MO_2_ (mass-specific values, no statistical analysis) was also plotted to visually explore trends within and between the offspring and parental acclimation groups (Fig. 5B). Overall, cold acclimated offspring attained a higher mean peak MO_2_ (mass-specific) than warm acclimated offspring (open vs. shaded boxes, W_♀_xW_♂_, C_♀_xW_♂_, W_♀_xC_♂_, C_♀_xC_♂_; Fig. 5B). When comparing offspring within an acclimation temperature, peak MO_2_ was comparable; cold acclimated offspring (open boxes, Fig. 5B) ranged between 335.8 ± 12.97 and 386.6 ± 13.13 mg O_2_ kg^−1^ h^−1^, and warm acclimated offspring (shaded boxes, Fig. 5B) ranged between 299.2 ± 18.79 and 325.5 ± 16.02 mg O_2_ kg^−1^ h^−1^.

### Critical thermal maximum (CTM)

Four AIC models best explained the trends in the CTM data (ΔAIC ≤ 2). Model 1 and 2 together suggest that CTM depended on a complex interaction between offspring (*T*_*O*_) and parental acclimation temperature (*T*_*M*_ and *T*_*F*_). The top model (W_i_ = 0.39) was the global model containing *Mass* as a covariate with the offspring (*T*_*O*_), maternal (*T*_*M*_) and paternal (*T*_*F*_) acclimation temperature as fixed effects, and all 2-way and 3-way interaction terms between them (Table 3). The second model was also the global model excluding *Mass*, however, model 1 was 1.40 (ER) times more likely to explain variation in CTM compared to model 2 (Table 3). The third model contained only *Mass* and offspring acclimation temperature (*T*_*O*_), whereas as the fourth model contained only *T*_*O*_ (Table 3).

Critical thermal maximum within and between the groups of offspring showed subtle differences (Fig. 5C). The mean CTM was comparable among groups of cold acclimated offspring (open boxes, Fig. 5C) with values ranging between 26.10 ± 0.2 and 26.64 ± 0.10 °C. Likewise, the CTM of warm acclimated offspring (shaded boxes, Fig. 5C) was similar. When comparing offspring within parental acclimation groups, warm acclimated offspring (shaded boxes) from W_♀_xW_♂_ parents had a CTM 1.17 °C higher than that of cold acclimated offspring (open boxes) from the same parental group (W_♀_xW_♂_, Fig. 5C). For the rest of the parental groups (C_♀_xW_♂_, W_♀_xC_♂_, C_♀_xC_♂_;), CTM was comparable (open vs. shaded boxes, Fig. 5C).

## Discussion

In this study, offspring from warm acclimated parents tolerated warmer temperatures better than those from cold acclimated parents, but only when the offspring’s environment was similar to the parental environment. By contrast, these offspring would be at a metabolic disadvantage at temperatures that are cooler than what their parents experienced. The results lend some support for the hypothesis that TGP occurs in cold-adapted, stenothermal ectotherms, potentially enabling them to cope with warmer environments. Although warm acclimation of the parents did not shift their offspring’s reaction norm upward as predicted, we found that the offspring’s thermal performance depended on complex interactions between parent and offspring environments. Ours is one of the few studies to investigate the relative parental contribution to TGP in a vertebrate offspring’s phenotype (Shama *et al*., 2014; Hellman et al,. 2019), and we demonstrate that the parents additively contribute to TGP.

### Evidence for transgenerational plasticity

With the acute temperature challenge, the MO_2_ of cold acclimated offspring from warm acclimated parents was elevated compared to those from cold acclimated parents at lower challenge temperatures (Fig. 2A), suggesting a higher cost of living (Norin & Metcalfe, 2019) at these temperatures when an environmental mismatch exists between generations. At warmer acute challenge temperatures (>18°C), cold acclimated offspring from the different mating crosses showed similar MO_2_ values, with declines beginning above 24°C (Fig. 2A). This disagrees with previous findings that warm acclimated offspring from warm acclimated parents had a lower metabolic rate (Donelson *et al*., 2012), although the effect of TGP can be difficult to predict and may not always be to the benefit of the offspring (Guillaume *et al*., 2016). In addition to the acute temperature challenge, the standard MO_2_ of offspring from warm acclimated parents (a generational environmental mismatch) was the highest among of the cold acclimated offspring (Fig. 5A), and parent acclimation temperature did not have an appreciable effect on peak MO_2_ (Fig. 5B). While this was in contrast to our predictions, standard metabolic rate is thought to be more plastic than maximum metabolic rate (Norin & Metcalfe, 2019) which could explain why standard MO_2_ was elevated in cold acclimated offspring from warm acclimated parents, whereas peak MO_2_ was relatively similar.

For the warm acclimated offspring, the acute thermal reaction norms were more similar regardless of the acclimation temperature of their parents (Fig. 2B). This was surprising given that previous studies have reported that offspring from warm acclimated parents could tolerate warm temperatures better than offspring from cold acclimated parents by reducing standard metabolic rate or increasing maximum metabolic rate (Donelson *et al*., 2012; Shama *et al*., 2014; Donelson *et al*., 2017). The disparity between the results of ours and earlier studies could be due to a number of reasons. First, these earlier studies tested TGP in tropical or eurythermal species, thus it is possible that TGP is limited in stenothermal species, like lake trout. Limited TGP may also be related to the limited variation in within-generation thermal plasticity in lake trout (Evans, 2007; Kelly *et al*., 2014). Also, because transgenerational effects (including epigenetic factors) are limited to available genetic resources (O’Dea *et al*., 2016; Donelson *et al*., 2017), it could be that lake trout simply do not have the capacity to extend their thermal tolerance (Evans, 2007; Kelly *et al*., 2014). Finally, it is possible that multiple generations of exposure to the same stressor may be required to strengthen the effect (Burggren, 2015; Bell & Hellman, 2019; Pilakouta et al., 2019), as in the case of the polychaete, *Ophryotrocha labronica*, where the effect of multigenerational exposure to warming was strongest in the F5 and F6 generations (Gibbin *et al*., 2017). Although a mechanistic explanation for the strengthening or weakening of a transgenerational effect has yet to be uncovered (Burggren, 2015), it may be that the effect of TGP on thermal tolerance and metabolic rate was relatively weak after only one generation of acclimation to warmer temperatures

Our results showed that the thermal experiences of the parents had a role in shaping the phenotype of the next generation, however, we did not explore the physiological mechanisms underlying variation in offspring metabolic rate. Nonetheless, TGP has been shown to act on mitochondrial capacity and gene expression (Shama *et al*., 2014; Gibbin *et al*., 2017). For example, mitochondria from the heart tissue of warm acclimated stickleback offspring from warm acclimated mothers had a lower rate of oxidative phosphorylation and less proton leak at warm temperatures (Shama *et al*., 2014). TGP has also been shown to up- or down-regulate the expression of genes involved in the heat shock response, metabolism, protein catabolism, immune response and reproduction (Veilleux *et al*., 2015; Shama *et al*., 2016; Veilleux *et al*., 2018). Mitochondrial function and gene expression would be useful to explore in lake trout to understand the mechanisms and limitations of TGP in this species and in cold-adapted, stenothermal organisms in general.

### Additive parental contribution

Transgenerational plasticity can be mediated through epigenetic modifications (summarized by Donkin & Barrès, 2018) that can be transmitted to the next generation (Crean & Bonduriansky, 2014; Marshall, 2015). Our results suggest that the contribution of the parents to TGP was additive. The offspring’s MO_2_ response to the acute temperature challenge was not influenced by the sole effect of either the maternal or paternal acclimation temperature, but instead on the complex interaction of maternal or paternal acclimation temperature with challenge temperature and offspring acclimation temperature (Table 2). The additive effect of parental temperatures on offspring metabolic rate was confirmed when both paternal and maternal acclimation temperature appeared in the top AIC models for standard and peak MO_2_ (Table 3). Both parents also contributed to their offspring’s upper thermal tolerance (CTM; Table 3) even though the differences in CTM among the groups of offspring was very slight (Fig. 5C). Similarly, Massamba-N’Siala et al. (2014) found no effect of TGP on the upper thermal tolerance of polychaetes given an acute temperature challenge.

In addition to epigenetic factors, other non-genetic effects (maternal and paternal) could have influenced the offspring’s phenotype (Burgess & Marshall, 2011; Shama, 2015). Females, for example, can provision their eggs through changes in egg size or nutrient enrichment of the yolk which can contribute to offspring fitness (Einum & Fleming, 1999; Gagliano & McCormick, 2007; Jonsson & Jonsson, 2016). To check for maternal provisioning in our study, we assessed egg size, mass and energy content, but did not find differences in these characteristics (non-significant result, data not shown) which agrees with Shama et al,. (2014) in that temperature acclimation of the parents did not affect egg size. Another maternal effect includes the transfer of hormones, such as thyroid or cortisol, to the eggs which could potentially alter offspring gene expression (Sopinka *et al*., 2017), growth and development (Gagliano & McCormick, 2009; Ruuskanen & Hsu, 2018). While we did not test the hormone content of the eggs, we acknowledge that it could potentially influence metabolic rate (Burton *et al*., 2011) of the offspring in our study.

The paternal contribution to TGP is understudied relative to maternal effects, although there is some evidence for non-genetic paternal effects (Crean & Bonduriansky, 2014; Marshall, 2015; Immler, 2018). Within the scope of TGP, the contribution of the father’s thermal environment to the offspring’s phenotype is variable with effects seen in some species (e.g. marine tubeworm: Guillaume *et al*., 2016) but not in others (e.g. stickleback: Shama *et al*., 2014). Further, the paternal contribution to TGP can extend beyond the transmission of epigenetic machinery to their offspring as ejaculate and sperm cytoplasmic components can also mediate paternal effects (Crean & Bonduriansky, 2014; Kekäläinen *et al*., 2018). It’s not entirely clear how these components could have affected the metabolic response of the offspring to temperature stress in our study.

### Perspectives and future directions

While TGP may have an important role in adaptation (Bernatchez, 2016; Smith *et al*., 2016), it is important to remember that some cold-adapted species have long generation times and TGP may not be able to keep up with the pace of climate change (Willi *et al*., 2006; Munday *et al*., 2013; Wilson *et al*., 2014). For example, lake trout can take 6 to 12 years to mature depending on latitude and lake productivity (Martin & Olver, 1980; Hansen *et al*., 2012). TGP could provide an advantage to populations with limited within-generation plasticity by buffering the negative effects of climate change between generations, thus giving the population a chance to adapt to elevated temperatures (Gibbin *et al*., 2017). Lake trout retreat to the cooler hypolimnion during the warmer summer months when the lake thermally stratifies (Casselman, 2008; Guzzo & Blanchfield, 2017), but climate change is expected to increase lake surface temperatures and prolong the duration of stratification (Lehman, 2002). For this reason, lake trout may be forced to reside in the hypolimnion for an extended period, lengthening their exposure to hypoxia which could negatively impact important life history traits (Evans, 2007; Guzzo & Blanchfield, 2017).

Other areas where an expanded understanding of the relationship between the environmental change and TGP is needed include the effect of multiple stressors (Gibbin *et al*., 2017; Karelitz *et al*., 2020), and the timing, strength and fluctuation of these stressors (Donelson *et al*., 2018). For example, few studies have examined the role of timing of an environmental perturbation with TGP in ectotherms (Massamba-N’Siala *et al*., 2014; Bell & Hellman, 2018). In our study, the parent lake trout were acclimated to a warm temperature only during the summer months when gametes were forming, but it would be interesting to see how an earlier onset of warming, or warming closer to breeding (typically mid-October for lake trout; Martin & Olver, 1980), affects the thermal performance of the offspring. It is not yet fully understood how the magnitude and fluctuation of stressors, such as extreme heat events, influences offspring thermal performance through TGP. Finally, it’s not yet clear how long transgenerational effects linger and whether these experiences can still be passed on to offspring long after the environmental perturbation subsides (reviewed in Herman *et al*., 2014).

From ours, and other studies (Yin *et al*., 2019), it is evident that TGP has a role to play in ‘priming’ offspring’s response to elevated temperatures. The physiological response to warming can include induction of heat shock proteins, changes in the concentration or isoform of proteins (e.g. enzymes and hormones), cellular membrane fluidity, and density of mitochondria (Horowitz, 2001; Lagerspetz, 2006; Bernabucci *et al*., 2010; Schulte, 2015). An understanding of how TGP acts to influence these physiological processes is warranted and will require further examination of the mechanisms underlying thermal tolerance, such as mitochondrial performance and gene expression in tandem with investigating which parental effects, including epigenetic inheritance (e.g. methylation, RNA interference), contribute to TGP.

We can conclude that TGP appears to occur in cold-adapted, stenothermal animals and may be an important factor for species having to cope with rapidly changing environmental conditions relating to climate change. An understanding of how phenotypic plasticity, developmental plasticity, TGP, and genetic changes combine to influence the adaptation of populations to climate change will not only help us anticipate the effects of a changing environment but will also deepen our knowledge of the link between plasticity, acclimation and adaptation.

## Acknowledgements

The Ontario Ministry of Natural Resources and Forestry (OMNRF) Fish Culture Section provided adult lake trout, rearing space and logistical support at the White Lake Fish Culture Station. Bill Sloan, Scott Ferguson, Anne McCarthy (OMNRF) and John Dewart provided assistance with breeding adult fish; Bill Sloan and Scott Ferguson also provided invaluable logistical support for all aspects of fish husbandry from care of the juvenile trout to assisting with experiments at the OMNRF Codrington Fisheries Research Facility. Caleigh Smith (OMNRF) genotyped adult and juvenile fish for parentage assignment and family identification of juveniles. Kurt Gamperl (Memorial University) provided advice on respirometer design, and Tim Johnson (OMNRF) provided access to a microcalorimeter to measure energy content in the eggs. Ben Bolker and John Dushoff (McMaster University) provided advice on statistical analysis.

