## Supplementary Info for "Limited transgenerational effects of environmental temperatures on thermal performance of a cold-adapted salmonid"

**DNA extraction and genotyping**

Genomic DNA of offspring lake trout was extracted from caudal fin samples, lysing approximately 0.25 $cm^2$ of tissue in deep-well 96-well plates by adding approximately 10 mg of tissue to each well along with 250 µL lysis buffer (50 mM Tris pH 8, 1000 mM NaCl, 1 mM EDTA, 1% sodium dodecyl sulphate (SDS) weight per volume, and 1000 µg proteinase K). The plates were incubated for 16 hours at 37°C, after which DNA was precipitated by adding 500 µL of 80% isopropanol per well and centrifuging the plates at 2000 *g* for 45 minutes. Afterwards, the supernatant was removed and the remaining pellets were rinsed with 1 mL of 70% ethanol, followed by re-centrifugation for 45 minutes at 2000 *g*. DNA pellets were air dried in a 70°C incubator for 30 minutes, then dissolved in 150 µl 1x TE (10 mM Tris, 1 mM EDTA). Extraction yields and quality were tested using electrophoresis alongside a mass ladder (Bioshop, Burlington, Ontario) in 1.5% agarose TBE gels stained with Sybr Green (Cedar Lane Laboratories, Burlington, Ontario).

Lake trout DNA samples were amplified at 17 microsatellite loci: MSU01, MSU02, MSU03, MSU05, MSU06, MSU08, MSU09, MSU10, MSU11, MSU13 (Rollins *et al*. 2009), *Ogo*1a (Olsen *et al*. 1998), *Sco*19 (Taylor *et al*. 2001), *Sco*215 (DeHaan *et al*., 2005), *Sfo*1, *Sfo*12 (Angers *et al*. 1995), *Sfo*C88 (King *et al*. 2012), and *Ssa*85 (O'Reilly *et al*. 1996). Multiplex reactions were performed in 10 µl reactions containing the following: 2 µl DNA with approximately 6 ng/µl, 1x PCR buffer containing 1.5 mM $MgCl_2$ (Qiagen, Mississauga, Ontario), 2 mM each dNTP (Bioshop, Burlington, Ontario), 0.5 mM $MgCl_2$ (Qiagen, Mississauga, Ontario), 0.2 mg/ml BSA (Bioshop, Burlington, Ontario), 0.025 U *Taq* DNA polymerase (Qiagen, Mississauga, Ontario) and dd$H_2O$. PCR cycling was carried out on Eppendorf Mastercycler Pro S thermal cyclers. Amplified products for all samples were run on an AB 3730 DNA analysis system with ROX 500 size standard (Applied Biosystems, Foster City, California). Allele sizes were scored using GeneMapper version 3.1 (Applied Biosystems, Foster City, California) and proofread with manual editing.

Sibling relationships were calculated for multilocus genotypes of the offspring using a maximum likelihood relatedness estimator in ML Relate (Kalinowski, Wagner & Taper, 2006). The breeding design (small number of parents, known closed mating history, and equal offspring family sizes) negated the need for more complex analytical approaches, and all offspring were assigned to specific mating crosses with high confidence.
